# N-cadherin orientational order decreases with mechanical load at cardiomyocyte adherens junctions

**DOI:** 10.64898/2026.07.17.739172

**Authors:** Yen T. B. Tran, William F. Dean, Yerin Han, Korina I. Karpov, Collin M. Ainslie, Alexa L. Mattheyses, Adam V. Kwiatkowski

## Abstract

Adherens junctions physically connect neighboring cells and are built around classical cadherins, homophilic transmembrane proteins that link to the actin cytoskeleton. Classical cadherins can organize into ordered arrays in vitro, but whether they do so in cells remains to be established. Here, we use fluorescence polarization microscopy to show that the classical cadherin N-cadherin is orientationally ordered at cardiomyocyte cell-cell junctions. Whereas the desmosomal cadherin desmoglein 2 was similarly ordered across junction types, N-cadherin order was spatially heterogeneous. Order was lowest where organized myofibrils terminate at high-load, vinculin-enriched axial junctions and highest at low-load, vinculin-poor lateral junctions. This inverse relationship between order and mechanical load suggests that robust cadherin-mediated adhesion does not require ectodomain order. Our findings provide evidence that a classical cadherin is orientationally ordered in cells and show that mechanically active adhesions adopt distinct organizational strategies according to local mechanical demands.

**Summary Statement:** At cardiomyocyte junctions, N-cadherin is ordered where mechanical load is low but disordered where load is high, suggesting that cadherin organization adapts to local force conditions.

## Introduction

Cell-cell adhesion is essential for tissue architecture and function. Adherens junctions (AJs) physically connect the actin cytoskeletons of neighboring cells (Campas et al., 2024). AJs are built around classical cadherins, transmembrane proteins that mediate homophilic adhesion, and are linked to the actin cytoskeleton through cytoplasmic catenin adaptor proteins (James et al., 2025; Troyanovsky, 2023). Structural analyses of purified type I cadherin ectodomains revealed that they can assemble into ordered arrays, leading to a model in which cadherins in AJs form a periodic two-dimensional lattice through lateral (*cis*) and adhesive (*trans*) interactions that promote adhesive strength (Harrison et al., 2011). Whether cadherins form such ordered assemblies within cells, and how any such organization is shaped by the cellular environment, is not known.

Classical cadherins are single-pass transmembrane proteins with five extracellular cadherin repeats (EC1-5). Opposing EC1 domains mediate Ca^2+^-dependent *trans* binding through a strand-swap mechanism, and adjacent cadherins engage in lateral *cis*-binding through an asymmetric EC1-EC2 interface (Boggon et al., 2002; Harrison et al., 2010; Shapiro et al., 1995; Wu et al., 2010; Wu et al., 2011). The intracellular cadherin tail binds p120-catenin and β-catenin, which in turn recruits the mechanosensitive F-actin-binding protein α-catenin (Campas et al., 2024; Charras and Yap, 2018; Mege and Ishiyama, 2017). Vinculin, another mechanosensitive actin-binding protein, is recruited under tension to further reinforce the cadherin-cytoskeletal connection (Huang et al., 2017; le Duc et al., 2010; Merkel et al., 2019; Pang et al., 2019; Yao et al., 2014; Yonemura et al., 2010). AJ adhesive strength and mechanical stability, therefore, depend on cadherin ectodomain *cis* and *trans* cadherin interactions, and cytoplasmic reinforcement of the cadherin-actin linkage. Studies in epithelial cells suggest that mechanical force promotes cadherin clustering and cytoplasmic reinforcement (Engl et al., 2014; le Duc et al., 2010; Leerberg et al., 2014; Strale et al., 2015; Truong Quang et al., 2013), but most examine AJs during initial formation or acute force application. It remains unclear how sustained, physiological mechanical load affects cadherin organization, and whether junctions under different loads adopt distinct organizational strategies.

Heart function requires the mechanical coupling of individual cardiomyocytes (CMs). Mature CMs are joined end-to-end by the intercalated disc, which comprises AJs and desmosomes linking the actin and intermediate filament cytoskeletons of adjoining cells (Vite and Radice, 2014). Immature CMs, by contrast, are apolar cells in which actomyosin filaments lack a dominant orientation and AJs instead form around the entire cell perimeter. As CMs mature and polarize, myofibrils reorient along a primary axis, and AJs segregate onto developing axial or lateral membranes (Granados-Riveron and Brook, 2012; Guo and Pu, 2020). This segregation creates two AJ populations in distinct mechanical environments within a single CM: axial AJs— where myofibrils terminate—bear high contractile forces, whereas lateral AJs running parallel to myofibrils experience relatively low load (Dong et al., 2025; Merkel et al., 2019; Vermij et al., 2017). Axial and lateral junctions also differ in the directionality of the force they sustain: axial junctions are predicted to bear primarily tensile force, whereas lateral AJs experience predominantly shear (Kale et al., 2018). Maturing CMs are therefore an ideal system to test how the magnitude and directionality of mechanical load relate to cadherin organization.

Here, we used fluorescence polarization microscopy (FPM) to investigate the nanoscale orientational order of cadherins at CM junctions. We focused on N-cadherin (neuronal cadherin, Ncad), the predominant classical cadherin at CM AJs (Kostetskii et al., 2005; Radice et al., 1997), and desmoglein 2 (Dsg2), a desmosomal cadherin expressed in CMs. We found that the ectodomains of both cadherins were ordered in CMs. While ectodomain order has been demonstrated for desmosomal cadherins in epithelial cells (Bartle et al., 2017; Dean and Mattheyses, 2022), our data provide evidence that order extends more broadly to CMs. Notably, to our knowledge, this study provides the first evidence that a classical cadherin is orientationally ordered at junctional membranes. Ncad order was heterogeneous, with the lowest order at high-load, vinculin-enriched axial junctions, and the highest order at low-load, vinculin-poor lateral junctions. We propose that mechanical load reduces Ncad order at axial junctions, creating an AJ architecture that can withstand high, multidirectional tension.

## Results

### Fluorescence polarization microscopy reveals N-cadherin and desmoglein 2 orientational order at cardiomyocyte cell-cell junctions

FPM quantifies protein orientational order within diffraction-limited regions and has revealed nanoscale protein organization in diverse cellular structures (Bartle et al., 2020; Bartle et al., 2017; Dean and Mattheyses, 2022; Dean and Mattheyses, 2024; Kampmann et al., 2011; Mehta et al., 2016). To probe cadherin order, we applied FPM to Ncad and Dsg2 in cultured mouse neonatal CMs. An FPM probe reporting Dsg2 extracellular domain orientation was previously validated (Dean and Mattheyses, 2022), but no such probe existed for Ncad. Therefore, we developed one, inserting EGFP between amino acids 314 and 315 in the EC2 domain (Ncad-EC2, Fig.1A) based on a published screen (Kim et al., 2011). Since EGFP is constrained by its N- and C-terminal associations in this construct (Fig. 1B), its orientation should reflect that of the EC2 domain. AlphaFold modeling predicted that the EC2 structure was unchanged in Ncad-EC2, with EGFP extending away from the *cis*-and *trans*-binding interfaces and not predicted to interfere with either (Fig. 1B, Fig. S1). Ncad with EGFP fused to the C-terminal tail (termed Ncad-C) was used as a control as the EGFP is unconstrained and should not correlate with Ncad orientation.

**Figure 1.**
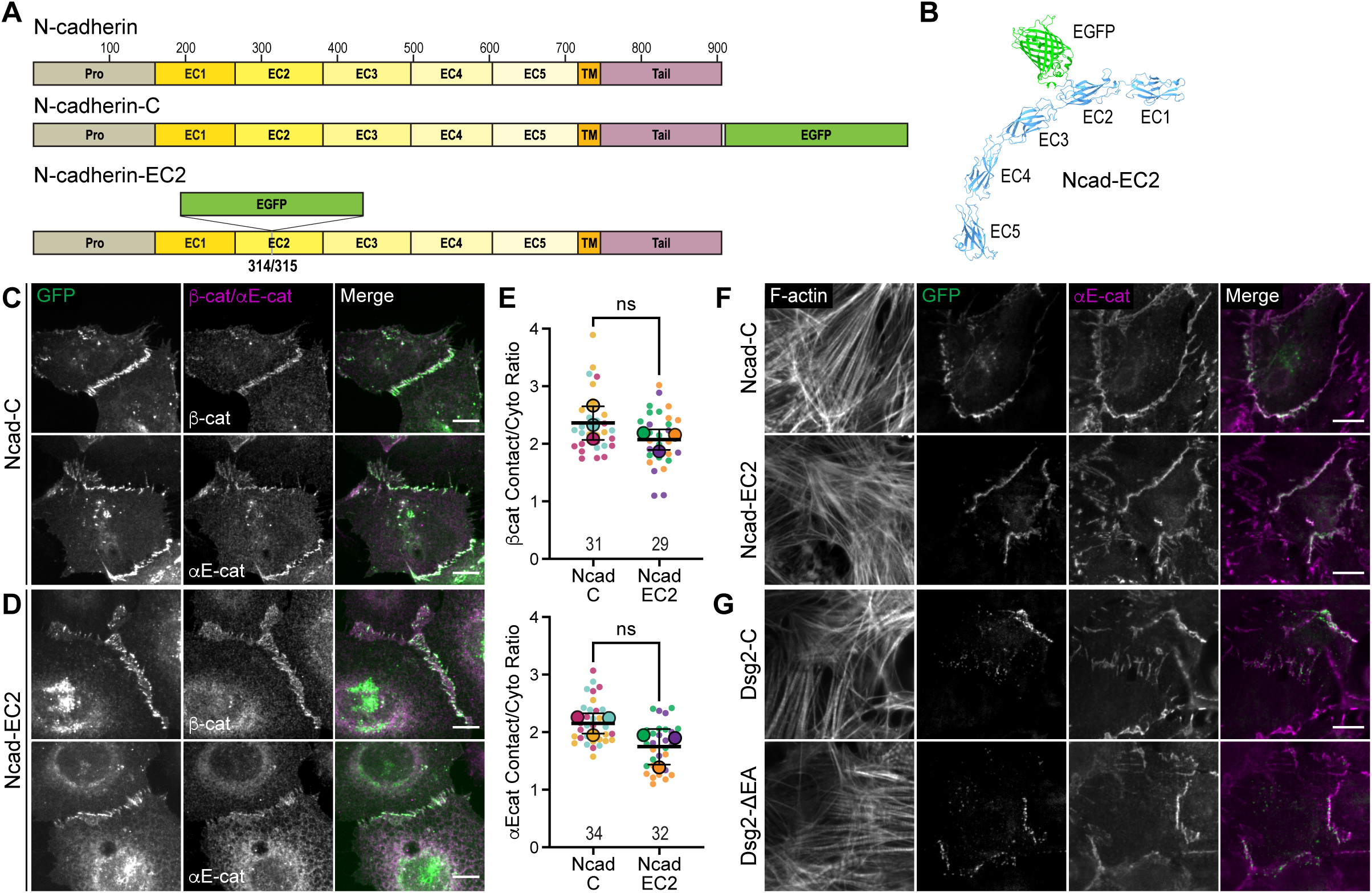
Ncad-EC2 reconstitutes epithelial AJs and localizes to cardiomyocyte AJs. (A) Schematic of N-cadherin domain organization and the EGFP-tagged constructs used in this study, with propeptide (PRO), extracellular cadherin (EC), transmembrane (TM), and tail regions marked. (B) AlphaFold model of the Ncad-EC2 extracellular domain. (C, D) Cadherin-deficient A431D cells expressing Ncad-C (C) or Ncad-EC2 (D), stained for endogenous β-catenin or αE-catenin. (E) Quantification of β-catenin (top) and αE-catenin (bottom) recruitment to cell-cell contacts (contact/cytoplasm ratio) in A431D cells expressing Ncad-C or Ncad-EC2. Data are presented as a SuperPlot (Lord et al., 2020). Error bars, mean ± SD; *n* (cells) below each column. Unpaired, two-tailed t-test; ns (not significant) p > 0.05. (F) GFP, αE-catenin, and F-actin in CMs expressing Ncad-C and Ncad-EC2 (G) or Dsg2-C and Dsg2-ΔEA. Scale bars, 10 µm (C, D, F, G).

First, we assessed whether Ncad-EC2 could form functional AJs. When expressed in cadherin-deficient A431D epithelial cells (Lewis et al., 1997), both Ncad-C and Ncad-EC2 restored cell-cell contact formation (Fig. 1C,D) and recruited β-catenin and αE-catenin at comparable levels (Fig. 1E), indicating that they form functionally similar AJs. Consistent with this, Ncad tagged at the same site with Venus was previously shown to reconstitute Ca^2+^-dependent adhesion in cadherin-deficient L cells (Kim et al., 2011). When expressed in mouse neonatal CMs, Ncad-EC2 localized to cell-cell junctions like Ncad-C (Fig. 1F; (Li et al., 2019; Merkel et al., 2019)). Thus, Ncad-EC2 is a functional cadherin that assembles AJs *de novo* and incorporates into existing CM AJs.

We also tested the localization of the established Dsg2 order probe, in which the extracellular anchor (EA) domain is replaced with EGFP (Dsg2-ΔEA) (Dean et al., 2024; Dean and Mattheyses, 2022). Like Ncad-C, the Dsg2 control has EGFP fused to the C-terminal tail (Dsg2-C). Both Dsg2 constructs localized to CM cell-cell junctions (Fig. 1G).

To measure order, we used excitation-resolved FPM, acquiring images at four distinct excitation polarizations (Fig. 2A). Fluorescence emission within a diffraction-limited spot contains signal from many individual EGFPs. If cadherins—and by extension, the constrained EGFPs—are in an ordered array, their EGFP absorption dipoles will be aligned and emission intensity will be sinusoidally modulated by excitation polarization. Conversely, if the cadherins are disordered, the EGFP absorption dipoles will be randomly oriented and fluorescence intensity will not vary systematically with excitation polarization The normalized modulation depth defines the order parameter, a dimensionless quantity ranging from zero (completely disordered) to one (fully ordered in the x,y plane) (Fig. 2B).

**Figure 2.**
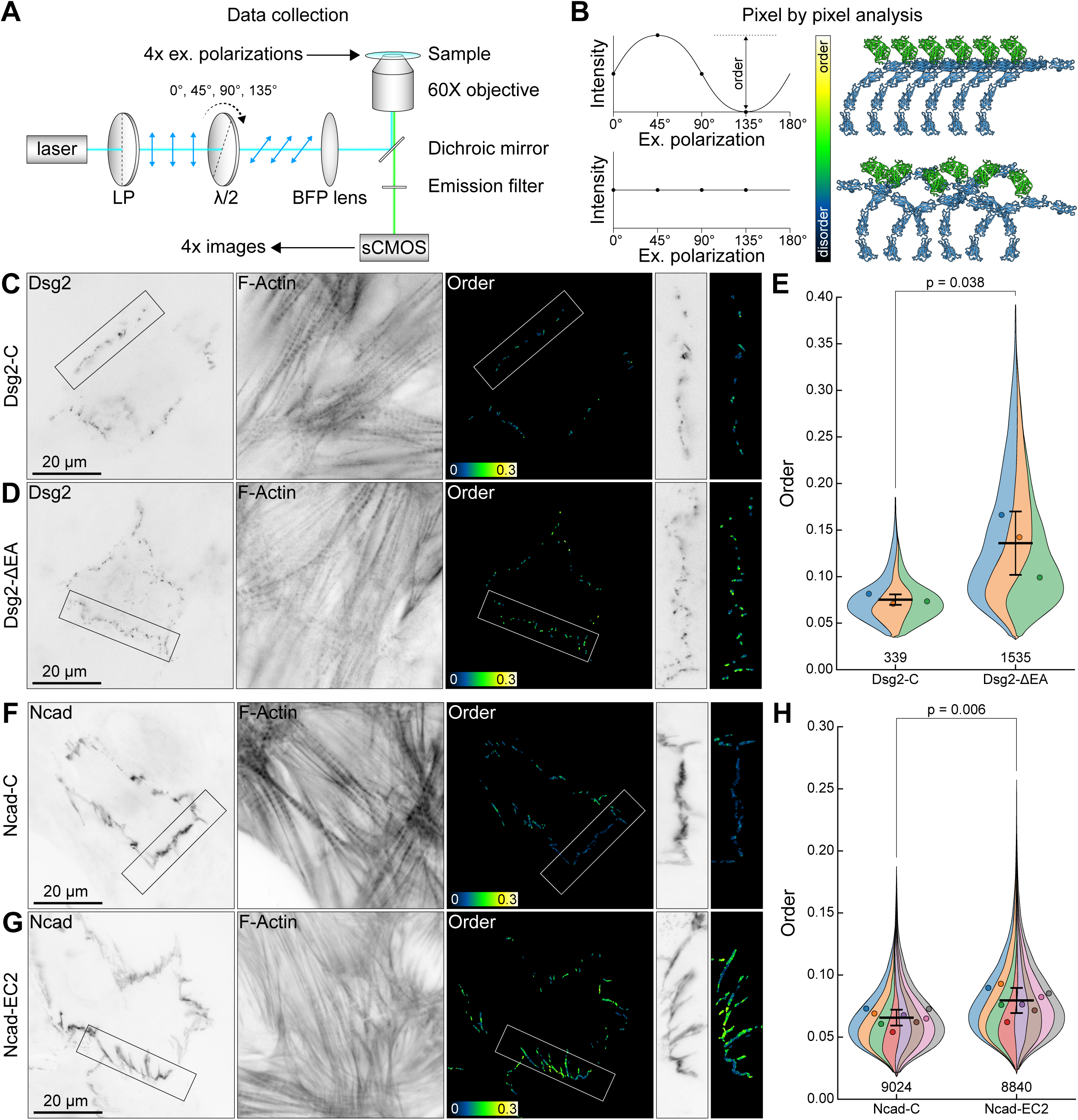
Ncad-EC2 is ordered in cardiomyocyte junctions. (A) Schematic of excitation-resolved FPM. A laser passes through a linear polarizer (LP) and a rotating half-wave plate (λ/2) before being focused on the objective back focal plane (BFP). Four images are acquired at excitation polarization angles 0°, 45°, 90°, and 135°. (B) If Ncad is ordered, Ncad-EC2 fluorescence intensity will be modulated by the excitation polarization. If Ncad is disordered, Ncad-EC2 emission will not vary with excitation polarization. (C, D) Dsg2 (EGFP signal), F-actin, and Dsg2 order in CMs expressing Dsg2-C (C) or Dsg2-ΔEA (D); rectangles mark the magnified cell-cell contacts at right. (E) Violin SuperPlot of Dsg2-C and Dsg2-ΔEA order. Colored stripes, biological replicates; circles, replicate means. (F, G) Ncad (EGFP signal), F-actin, and Ncad order in CMs expressing Ncad-C (F) or Ncad-EC2 (G). (H) Violin SuperPlot of Ncad-C and Ncad-EC2 order. Error bars, mean ± SD of the biological replicates; unpaired two-tailed t-test (E, H). Scale bars, 20 µm.

We first asked whether Dsg2 is ordered at CM junctions. The Dsg2-C control displayed low order (0.075 ± 0.006), whereas Dsg2-ΔEA was significantly more ordered (0.134 ± 0.034, unpaired t-test, p=0.038; Fig. 2C-E). We conclude that Dsg2 ectodomains are ordered at CM junctions, similar to Dsg2, Dsg3, and desmocollin 2 (Dsc2) ectodomains in epithelial cells (Bartle et al., 2017; Dean et al., 2024; Dean and Mattheyses, 2022).

We then examined Ncad order. As expected, the control Ncad-C showed low order (0.066 ± 0.006; Fig. 2F,H). In contrast, Ncad-EC2 was significantly more ordered (0.080 ± 0.010, unpaired t-test p=0.006; Fig. 2G,H), indicating that the Ncad ectodomain is ordered at CM junctions. Because FPM reports the orientation of the tagged domain—the EC2 domain for Ncad and the membrane-proximal EA domain for Dsg2—order values are not directly comparable between the two probes. Notably, the level of order observed in Ncad-EC2 suggests that Ncad does not organize into an ideal crystalline lattice. Instead, the data suggest a preference for a particular orientation and could represent a loosely packed, dynamic array. Nonetheless, these results demonstrate that Ncad, like its desmosomal cadherin counterparts, can achieve orientational order at junctional membranes.

### N-cadherin is more ordered at lateral than at axial junctions

While reviewing the order images, we noted a striking difference between the two cadherins: Dsg2 order was relatively uniform (Fig. 2D), whereas Ncad order was markedly heterogeneous (Fig. 2G). Neonatal CMs remodel junctions in culture (Li et al., 2019). Within two to three days of culture, two junction types begin to emerge as CMs polarize: axial junctions where myofibrils terminate, and lateral junctions that run parallel to myofibrils (Fig. 3A,B). Axial junctions experience high loads and are enriched for vinculin, whereas lateral junctions experience lower loads and have limited vinculin.

**Figure 3.**
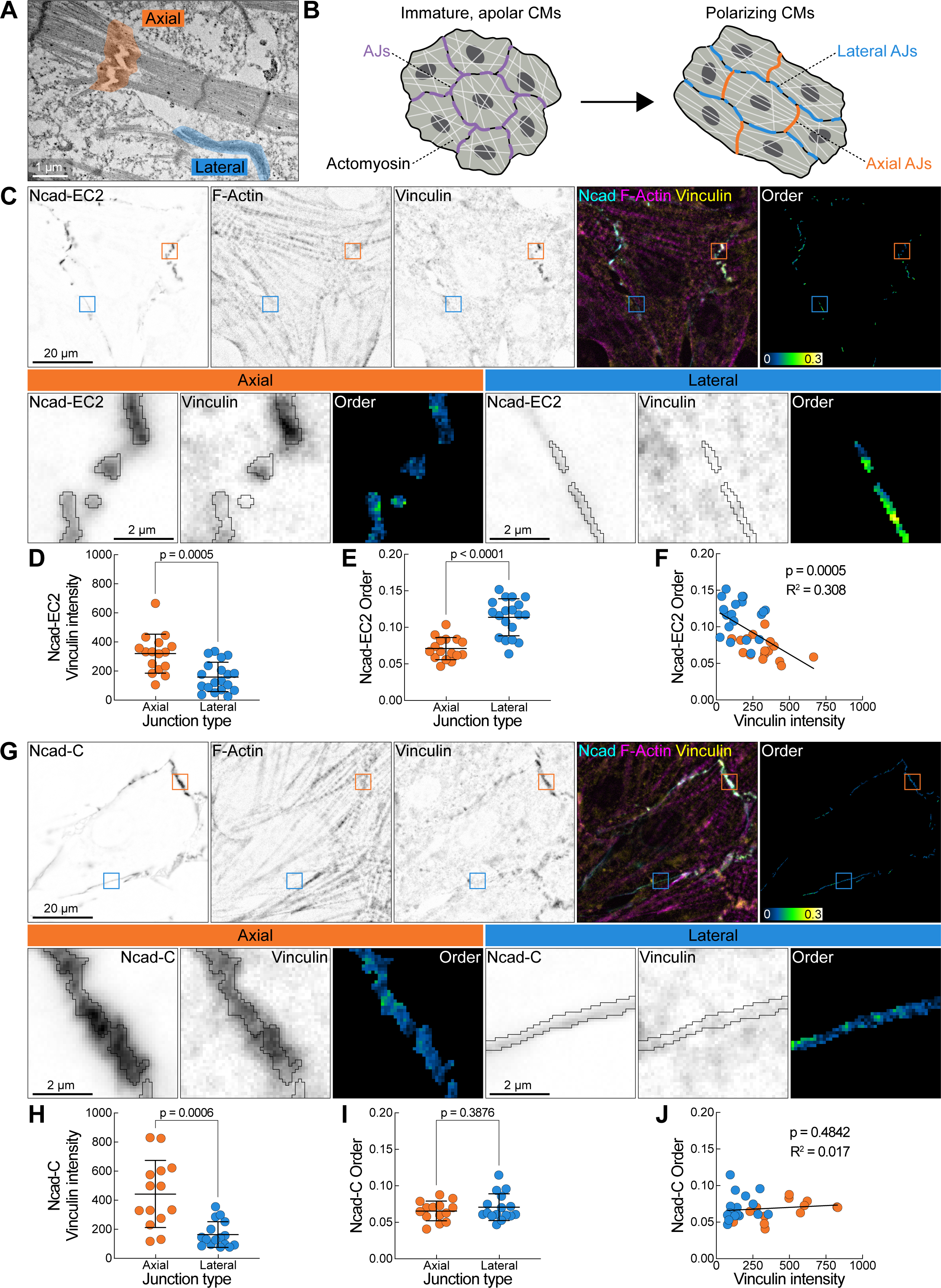
Ncad-EC2 is less ordered in axial junctions. (A) Electron micrograph of cell-cell contact between cultured neonatal mouse CMs; axial and lateral junctions in orange and blue, respectively. (B) Junction organization in CMs. Immature CMs are apolar cells encircled by AJs; as CMs polarize, AJs segregate onto axial (orange) or lateral (blue) membranes as myofibrils reorient and organize. (C) CM expressing Ncad-EC2 stained for F-actin and vinculin, with merge (Ncad, cyan; F-actin, magenta; vinculin, yellow) and order image. Axial and lateral junctions (color-coded squares) identified by actin organization. Bottom panels, magnified axial and lateral junctions. (D, E) Vinculin intensity (D) and Ncad-EC2 order (E) in axial and lateral junctions in Ncad-EC2 expressing CMs. (F) Ncad-EC2 order versus vinculin intensity in axial (orange) and lateral (blue) junctions. (G) CM expressing Ncad-C, labeled and imaged as in (C). (H, I) Vinculin intensity (H) and Ncad-C order (I) in axial and lateral junctions in Ncad-C expressing CMs. (J) Ncad-C order versus vinculin intensity in axial (orange) and lateral (blue) junctions. Error bars, mean ± SD, unpaired two-tailed t-test. (E) Ncad-EC2 order in axial and lateral junctions. Mean ± SD, unpaired two-tailed t-test (D, E, H, I). Slope and R^2^ from linear regression analysis; slope deviation from zero analyzed for significance (F, J). Data in D-F from 16 axial (3 outliers removed) and 19 lateral ROIs; data in H-J from 14 axial (3 outliers removed) and 16 lateral (1 outlier removed) ROIs. Scale bars, 1 µm (A), 20 µm, 2 µm (C, G, insets).

To determine whether Ncad order differed with junction type, we analyzed axial or lateral regions of interest (ROI) spanning multiple junctions in CMs expressing Ncad-EC2 (Fig. 3C). As expected, vinculin intensity was higher in axial regions than in lateral regions (axial, 320.6 ± 134.2; lateral, 159.9 ± 100.0; Fig. 3D). Strikingly, Ncad-EC2 order was significantly higher at lateral junctions (0.114 ± 0.025) than at axial junctions (0.071 ± 0.015, Fig. 3E), and Ncad-EC2 lateral order was higher than the Ncad-C baseline (0.066 ± 0.006; Fig. 2H), indicating that Ncad forms ordered arrays at lateral junctions. Across both axial and lateral regions, vinculin intensity and order were inversely correlated (Fig. 3F). Repeating the analysis in CMs expressing Ncad-C (Fig. 3G), vinculin was again significantly enriched in axial regions (axial, 443.0 ± 230.6; lateral, 163.9 ± 88.4; Fig. 3H), but order did not differ between regions (axial, 0.066 ± 0.013; lateral, 0.071 ± 0.018; Fig. 3I). Consistent with this, no correlation was observed between vinculin intensity and order in Ncad-C (Fig. 3J). Notably, Ncad-EC2 order at axial junctions was indistinguishable from Ncad-C (Fig. 3E,I), indicating that Ncad order was lost at axial junctions. We conclude that Ncad order is higher at lateral junctions than at axial junctions, suggesting that either the magnitude or direction of load disrupts Ncad order.

### N-cadherin order is inversely correlated with vinculin enrichment

Beyond the difference between axial and lateral junctions, we observed striking variability in Ncad order within individual junctions (Fig. 4A,B). We asked whether the variability in Ncad order reflected differences in load across individual junctions, again using vinculin enrichment as a proxy for load (Kale et al., 2018). We postulated that mixed-order junctions contain a gradient of vinculin recruitment inversely correlated with order. Indeed, mixed junctions contained adjacent regions of high and low vinculin intensity, and linescan analysis confirmed that order was inversely correlated with vinculin within individual junctions (Fig. 4C).

**Figure 4.**
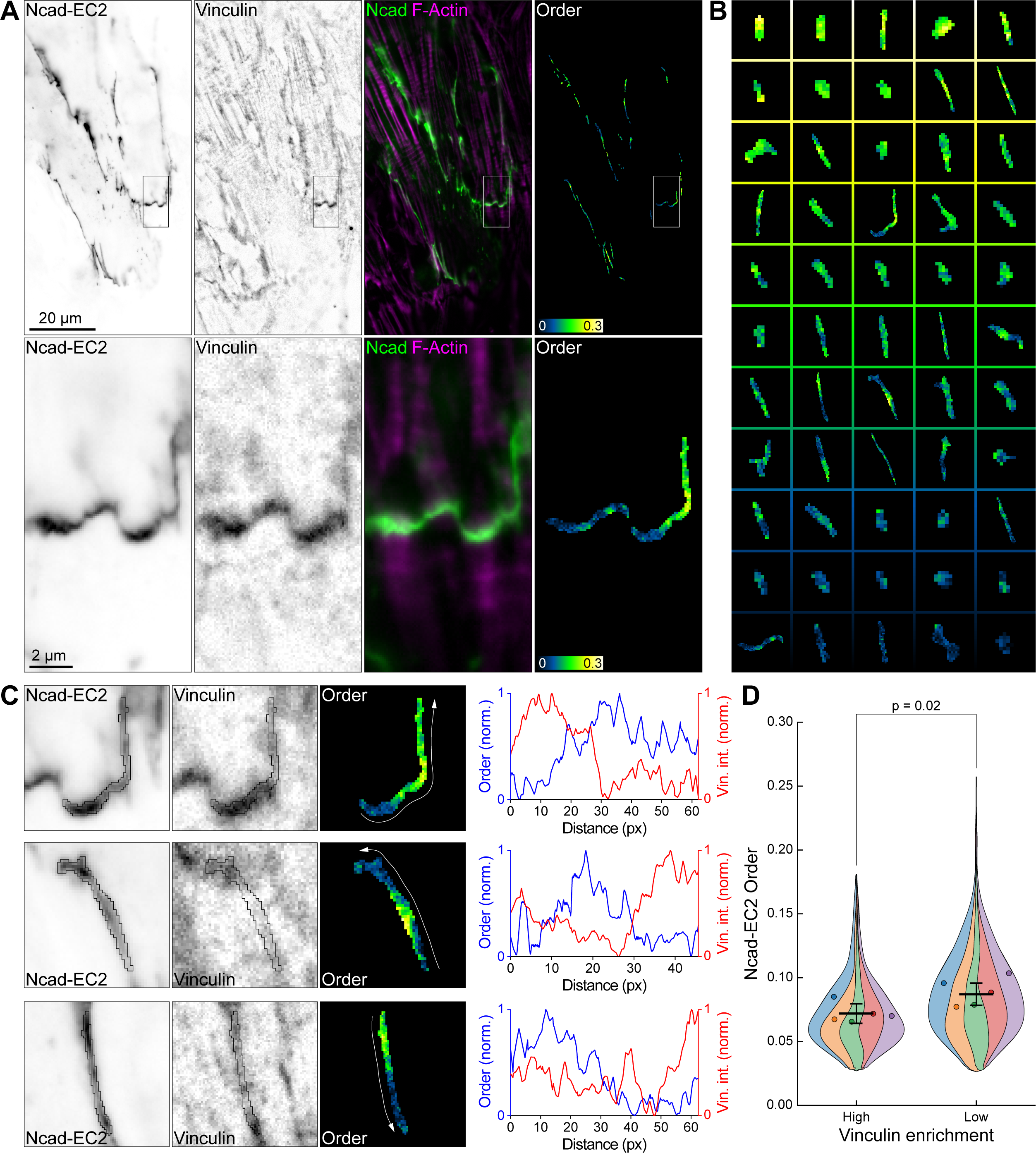
Ncad-EC2 order is heterogeneous within individual junctions. (A) CM expressing Ncad-EC2 stained for vinculin, with merge (Ncad, green; F-actin, magenta), and order image. (B) Ncad-EC2 order in individual junction objects from the cell in (A), showing heterogeneous order. (C) Magnified Ncad-EC2, vinculin, and order images from three junctions in (B), with linescans of normalized order (blue) and vinculin intensity (red) along each junction. White arrows mark linescan direction. (G) Violin SuperPlot of Ncad-EC2 order in junctions with high (top 10%) versus low (bottom 10%) vinculin intensity. Error bars, mean ± SD; unpaired two-tailed t-test. Scale bars, 20 µm, 2 µm (A, inset).

We next asked whether the inverse relationship between vinculin enrichment and order held across junctions. For each Ncad-EC2 experiment, we identified junctions in the top and bottom 10% of vinculin intensity and then compared the mean order between the groups. Junctions with the highest vinculin intensity had significantly less Ncad-EC2 order than those with the lowest (0.072 ± 0.008 vs. 0.087 ± 0.009; Fig. 4D), supporting the conclusion that Ncad is less ordered in vinculin-enriched junctions.

## Discussion

Our results show that AJs adopt different organizational strategies at low- and high-load junctions. This difference could reflect either of two coupled features of the load environment: magnitude and directionality. Axial junctions couple contractile myofibrils between cells and bear the brunt of repeated actomyosin contraction, experiencing substantially higher force than lateral junctions, which run largely parallel to the contractile filaments. Axial and lateral junctions also differ in load directionality: based on their orientation relative to the contractile axis, axial junctions bear primarily tensile force, whereas lateral junctions predominantly experience shear. As load magnitude and directionality covary across these populations, we consider each in turn.

Ncad order may be tuned to load directionality. At lateral junctions, an ordered cadherin array would distribute shear forces roughly equally across cadherin-catenin complexes in similar geometric environments. Such an array could also be remodeled through coordinated reorientation of *cis* interactions rather than breaking *trans* bonds. By accommodating shear rather than resisting it stiffly, an ordered cadherin array could provide a non-destructive deformation pathway that maintains lateral junctions during contraction. At axial junctions, by contrast, Ncad disorder (cadherin-catenin complexes in many different orientations) would be well-suited for bearing multidirectional tensile load. A disordered network with no preferred orientation has no single direction along which it is weakest. Because the complexes adopt varied local geometries, individual *trans* bonds bear different shares of the load and would fail individually across a range of cellular loads rather than all together under a critical load. Such an isotropic, disordered network is well-suited for variable, multidirectional tension that arises as developing myofibrils integrate into axial junctions.

The loss of Ncad order at axial junctions could instead reflect the higher magnitude of load they bear. In this model, increased load recruits cytoplasmic force-coupling adaptor proteins. Structural modeling suggests that a cadherin lattice sterically excludes vinculin (Troyanovsky et al., 2021). Because vinculin recruitment scales with force (Kale et al., 2018; Seddiki et al., 2018), junctions under higher load would be expected to lose ectodomain order as these complexes are built. This is consistent with the inverse correlation we observed between vinculin recruitment and Ncad order. High-load junctions may therefore favor cytoplasmic reinforcement at the expense of ordered ectodomain arrays. We note that most junctional complexes, both axial and lateral, were heterogeneous, containing a mix of ordered and disordered regions rather than a single uniform state. This indicates order is regulated locally within junctions and suggests that load magnitude and directionality likely act together to tune cadherin order. Resolving the interplay between cadherin ectodomain organization, load magnitude, force direction, and adhesion strength will require further work.

Two further aspects of the relationship between order and junction function are notable. First, robust cadherin-mediated adhesion does not require ectodomain order: axial junctions, which couple contractile myofibrils and must sustain adhesion under strong loads, were the least ordered. Such decoupling of cadherin order and adhesion was observed in hyperadhesive desmosomes, where removal of calcium caused Dsg3 to become disordered, but adhesion was maintained (Bartle et al., 2017). Second, our data indicate that cadherin order is not templated by cytoskeletal order. Highly organized myofibrils terminate at axial junctions, yet these junctions showed the lowest Ncad order. Cytoskeletal organization thus does not dictate classical cadherin order, at least during active junction remodeling, but could once CMs complete polarization.

### Limitations

The FPM resolution is diffraction-limited and cannot resolve cadherin nanoclusters. Individual nanoclusters could contain ordered cadherins, but a collection of nanoclusters in an AJ without a particular orientation between them would appear disordered in our current imaging system. In addition, the more complex topology of axial junctions could, in principle, confound order analysis, leading to a decrease in observed order. However, low-order, high-vinculin regions were occasionally observed at lateral junctions, indicating that order tracks vinculin enrichment rather than junction geometry.

### Conclusion

This study reveals how cadherin nanoscale organization varies with mechanical load in a cellular context. We show that Ncad order is not required for robust adhesion and that junctions adopt distinct organizational strategies based on mechanical context. The inverse relationship between Ncad order and vinculin recruitment suggests that junctions do not simply recruit more proteins under load, but additionally reorganize their architecture, trading ordered ectodomain arrays for cytoplasmic reinforcement to strengthen adhesion. We speculate that this load-dependent reorganization enables dynamic remodeling of adhesion during myofibril growth and coupling—processes essential for cardiac development and adaptation.

## Materials and Methods

### Constructs

Mouse N-cadherin in pEGFP-N1 (Ncad-EGFP-C) was a gift from James Nelson (Stanford University). To make Ncad-EC2, the N-cadherin CDS was amplified from Ncad-C and cloned into pcDNA3.1. Next, EGFP was inserted between N-cadherin amino acids 314 and 315 by Gibson Assembly to create Ncad-EC2. Dsg2 constructs were previously published with Dsg2-ΔEA as Dsg2-ECTO and Dsg2-C as Dsg2-LINK (Dean and Mattheyses, 2022).

### Cardiomyocyte isolation and culture

All animal work was approved by the University of Pittsburgh Animal Research Protection Office. Primary CMs were isolated from postnatal day 1 Swiss Webster mice of both sexes as described previously (Ehler et al., 2013). CMs were plated onto 35 mm MatTek dishes with 10 mm insets coated with Collagen Type I (Millipore). CMs were plated in plating media: 65% DMEM (Thermo Fisher Scientific), 19% M-199 (Thermo Fisher Scientific), 10% horse serum (Thermo Fisher Scientific), 5% fetal bovine serum (Atlanta Biologicals) and 1% penicillin-streptomycin (Thermo Fisher Scientific). Media was replaced 16 hours after plating with maintenance media: 78% DMEM, 17% M-199, 4% horse serum, 1% penicillin-streptomyocin, 1 µM AraC (Sigma), and 1 µM Isoproterenol (Sigma). CMs were cultured in maintenance media until fixation.

### Cell culture and transfection

Cadherin-deficient A431D cells (Lewis et al., 1997) were cultured in DMEM plus 10% fetal bovine serum, 1% penicillin-streptomycin. CMs and A431D cells were transfected with Lipofectamine 2000 (Thermo Fisher Scientific) one day after plating.

### Cell fixation and immunostaining

All cells were fixed in warmed (37°C) 4% EM grade paraformaldehyde in PBS (Ca^2+^, Mg^2+^) plus 120 mM sucrose for 10 min at room temperature. Cells were washed twice with PBS (Ca^2+^, Mg^2+^). Cells were permeabilized with 0.2% Triton X-100 in PBS (Ca^2+^, Mg^2+^) for 5 min and washed twice with PBS (Ca^2+^, Mg^2+^). Cells were blocked in 10% bovine serum albumin (BSA; Sigma) in PBS (Ca^2+^, Mg^2+^) for 1 hr at room temperature. Samples were incubated with primary antibodies in PBS (Ca^2+^, Mg^2+^) + 1% BSA for 1 hr at room temperature, washed twice in PBS, incubated with secondary antibodies in PBS (Ca^2+^, Mg^2+^) + 1% BSA for 1 hr at room temperature, washed twice in PBS (Ca^2+^, Mg^2+^), and then stored in PBS (Ca^2+^, Mg^2+^) or mounted in Prolong Glass (Thermo Fisher Scientific). All mounted samples were cured for at least 24 hr before imaging.

### Antibodies

Primary antibodies used for immunostaining: anti-αE-catenin (1:100, Enzo Life Science, ALX-804-101-C100), anti-β-Catenin (1:100, Cell Signaling, D10A8), anti-vinculin (1:800; Sigma Aldrich V9131). F-actin was visualized using an Alexa Fluor dye conjugated to phalloidin (1:100; Thermo Fisher Scientific). Secondary antibodies used were goat anti-mouse conjugated to Alexa Fluor dyes (1:250, Thermo Fisher Scientific).

### Confocal microscopy imaging and analysis

Transfected A431D and CMs in Fig. 1 were imaged with a 100x NA 1.45 oil-immersion objective on a Nikon Eclipse Ti inverted microscope outfitted with a Prairie swept field confocal scanner, Agilent monolithic laser launch and Andor iXon3 camera using NIS-Elements (Nikon) imaging software. All other images were acquired following the *Fluorescence Polarization Microscopy* protocol described below.

To quantify β-catenin and α-catenin recruitment to cell-cell contacts in A431D cells expressing Ncad-C or Ncad-EC2, transfected cells were fixed, stained for β-catenin or αE-catenin, and imaged. For analysis, maximum projections of Z-stacks spanning 1 μm (six sections at 200 nm intervals) were created, centered on the junction focal plane. In the maximum projection image, the GFP channel was thresholded to create a binary mask of the cell-cell contacts. Mean β-catenin or αE-catenin fluorescence intensity was measured within the cell-cell contact mask and in three cytoplasmic regions within the transfected cell. To calculate the contact/cytoplasm ratio, the β-catenin or αE-catenin fluorescence intensity within the masked region was divided by the average cytoplasmic signal to normalize between samples. Data were plotted as SuperPlots (Lord et al., 2020) in Prism software (GraphPad).

### Fluorescence Polarization Microscopy

CMs were transfected with FPM probes and cultured for two days before fixation. Cells were stained and then imaged using FPM as previously described (Dean and Mattheyses, 2022). Briefly, cells were imaged with a Nikon Ti2 microscope equipped with a motorized stage and a 60x NA 1.49 oil-immersion objective. A 488 nm laser (Coherent) operating at 15 mW was directed through a linear polarizer and an achromatic half-wave plate (Thorlabs) before being focused at the back focal plane of the objective via a focusing lens. The half-wave plate was mounted on a motorized rotation stage (Thorlabs) to control the excitation polarization angle. Fluorescence excitation and emission were passed through a dichroic cube (89902 ET-405/488/561/647nm) and an additional bandpass emission filter (ET525/50m, Chroma). Images were recorded with an ORCA-Flash 4.0 v3 CMOS camera (Hamamatsu). For each field of view, a polarization-resolved image stack was acquired, with images at 4 excitation angles (0°, 45°, 90°, and 135°), each with an exposure time of 50 ms. To correct for spatial and polarization-dependent variations in illumination, flat-field calibration images were acquired prior to each experiment using an auto-fluorescent plastic slide (92001; Chroma). Five calibration stacks collected at different x-y positions were averaged at each polarization angle. The averaged calibration stack was normalized to its maximum intensity to generate the correction profile applied during image processing.

### Epifluorescence microscopy

After acquiring the FPM image, epifluorescence images were acquired using a Sola excitation light source with 488 (Filter cube: 96362, ET GFP, C180702), 561 (Filter cube: 96365 ET/mCH/TR C180338), and 647 (Filter cube: 96365 ET CY5 C180340) channels, as appropriate for the samples. Imaging was performed sequentially on the same microscope as FPM.

### Fluorescence Polarization Microscopy Image Analysis

Images were analyzed in MATLAB (The MathWorks, Natick, MA) using object-oriented polarization software (OOPS) (Dean et al., 2024b). Raw fluorescence polarization datasets were first flat-field corrected for nonuniform illumination and polarization-dependent variations in excitation intensity. To calculate the order (*p*) in each image pixel, a formulation based on the first three Stokes parameters was used:

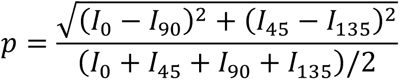

Where *I*_0_, *I*_45_, *I*_90_, and *I*_135_ are the flat-field-corrected emission intensities for each excitation polarization angle.

### Junction identification

Images were segmented into object-based masks based on local EGFP intensity using Otsu’s method. Objects located in the central part of the cell, and not at a cell-cell junctions, were excluded from the analysis. Junctions between two transfected cells were also removed to eliminate the possibility of homoFRET from *trans*-binding. The resulting masks were applied to the order images for quantification. The mean order within each connected component (junction) was calculated (Figs. 2,4). In Fig. 3, ROIs were defined based on the presence or absence of F-Actin attachments to the EGFP-tagged Ncad constructs. Within each ROI, the mean order of the masked pixels in the ROI was calculated. These same regions were applied to the vinculin intensity channel. Outliers were identified separately for axial and lateral junctions using the Tukey 1.5×IQR rule. For each junction type, values below Q1 − 1.5×IQR or above Q3 + 1.5×IQR were flagged as outliers and excluded from the analysis. In Fig. 4, the mean order from junctions with vinculin intensity less than or equal to the 10th percentile cutoff and junctions with vinculin intensity greater than or equal to the 90th percentile cutoff within individual experiments were compared. All vinculin images were preprocessed with a rolling ball background subtraction in Fiji prior to analysis.

All order images are displayed using the “GreenFireBlue” lookup table from Fiji (blue = low order, green = moderate order, and yellow = high order. Data were visualized as Violin SuperPlots using the superviolin Python package (Kenny and Schoen, 2021), which overlays the pooled single-measurement distribution with the per-replicate means, color-coding measurements by biological replicate. Error bars show the replicate mean ± SD.

## Acknowledgements

We are grateful to the Mattheyses and Kwiatkowski labs for generous and thoughtful discussion throughout this work.

## Competing interests

No competing interests declared.

## Funding

This work was supported by National Institutes of Health R01 AR072697 to ALM and R01 HL127711 to AVK.

---

**Fig. S1.**
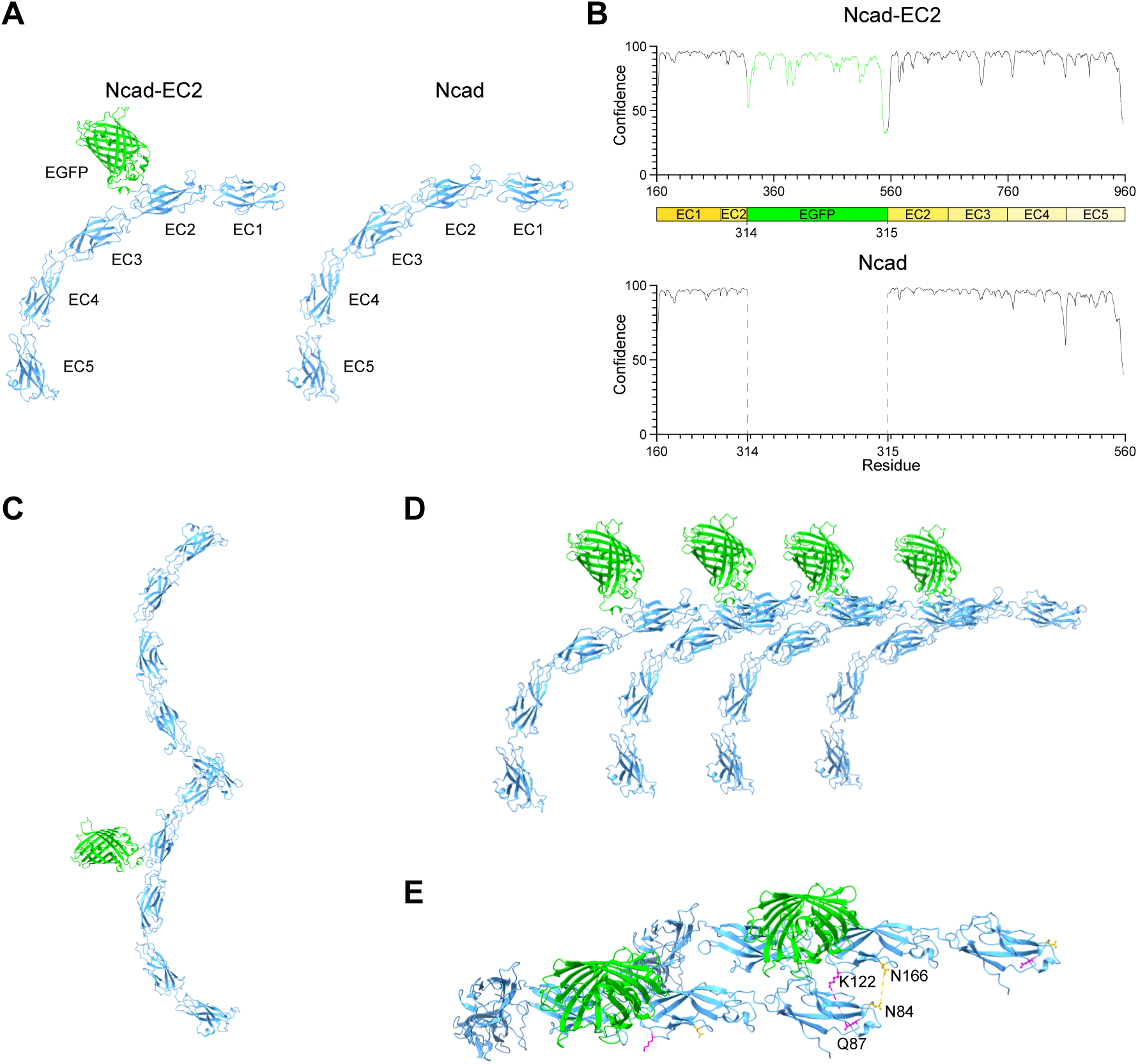
Ncad-EC2 predicted architectures. (A) AlphaFold predictions of the extracellular domains of Ncad-EC2 and wildtype (WT) NCad with EC domains labeled. (B) Confidence per residue plots for the models in A. (C) Ncad-EC2 and WT Ncad in a *trans*-binding conformation shows EGFP extending away from the binding site. (D, E) Ncad-EC2 undergoing *cis*-interactions viewed from the side (D) or the top down showing the location of key interacting residues (E).

## Notes

### Competing Interest Statement

The authors have declared no competing interest.

## References

Bartle, E.I., T.C. Rao, R.R. Beggs, W.F. Dean, T.M. Urner, A.P. Kowalczyk, and A.L. Mattheyses. 2020. Protein exchange is reduced in calcium-independent epithelial junctions. J Cell Biol. 219.

Bartle, E.I., T.M. Urner, S.S. Raju, and A.L. Mattheyses. 2017. Desmoglein 3 Order and Dynamics in Desmosomes Determined by Fluorescence Polarization Microscopy. Biophysical journal. 113:2519–2529.

Boggon, T.J., J. Murray, S. Chappuis-Flament, E. Wong, B.M. Gumbiner, and L. Shapiro. 2002. C-cadherin ectodomain structure and implications for cell adhesion mechanisms. Science. 296:1308–1313.

Campas, O., I. Noordstra, and A.S. Yap. 2024. Adherens junctions as molecular regulators of emergent tissue mechanics. Nature reviews. Molecular cell biology. 25:252–269.

Charras, G., and A.S. Yap. 2018. Tensile Forces and Mechanotransduction at Cell-Cell Junctions. Curr Biol. 28:R445–R457.

Dean, W.F., R.M. Albert, T.J. Nawara, M. Ubil, R.R. Beggs, and A.L. Mattheyses. 2024. Dsg2 ectodomain organization increases throughout desmosome assembly. Cell adhesion & migration. 18:1–13.

Dean, W.F., and A.L. Mattheyses. 2022. Defining domain-specific orientational order in the desmosomal cadherins. Biophysical journal. 121:4325–4341.

Dean, W.F., and A.L. Mattheyses. 2024. Illuminating cellular architecture and dynamics with fluorescence polarization microscopy. J Cell Sci. 137.

Dong, Y., A. Elgerbi, B. Xie, Y. Han, A.V. Kwiatkowski, J.S. Choy, and S. Sivasankar. 2025. Actomyosin forces trigger a conformational change in desmoplakin within desmosomes. Nat Commun. 16:9052.

Ehler, E., T. Moore-Morris, and S. Lange. 2013. Isolation and culture of neonatal mouse cardiomyocytes. J Vis Exp.

Engl, W., B. Arasi, L.L. Yap, J.P. Thiery, and V. Viasnoff. 2014. Actin dynamics modulate mechanosensitive immobilization of E-cadherin at adherens junctions. Nat Cell Biol. 16:587–594.

Granados-Riveron, J.T., and J.D. Brook. 2012. The impact of mechanical forces in heart morphogenesis. Circulation. Cardiovascular genetics. 5:132–142.

Guo, Y., and W.T. Pu. 2020. Cardiomyocyte Maturation: New Phase in Development. Circulation research. 126:1086–1106.

Harrison, O.J., F. Bahna, P.S. Katsamba, X. Jin, J. Brasch, J. Vendome, G. Ahlsen, K.J. Carroll, S.R. Price, B. Honig, and L. Shapiro. 2010. Two-step adhesive binding by classical cadherins. Nat Struct Mol Biol. 17:348–357.

Harrison, O.J., X. Jin, S. Hong, F. Bahna, G. Ahlsen, J. Brasch, Y. Wu, J. Vendome, K. Felsovalyi, C.M. Hampton, R.B. Troyanovsky, A. Ben-Shaul, J. Frank, S.M. Troyanovsky, L. Shapiro, and B. Honig. 2011. The extracellular architecture of adherens junctions revealed by crystal structures of type I cadherins. Structure. 19:244–256.

Huang, D.L., N.A. Bax, C.D. Buckley, W.I. Weis, and A.R. Dunn. 2017. Vinculin forms a directionally asymmetric catch bond with F-actin. Science. 357:703–706.

James, J., L.B.A. Winn, P. Mottram-Epson, and D. Koster. 2025. Paths to stability - actin regulation of adherens junction mechanics. J Cell Sci. 138.

Kale, G.R., X. Yang, J.M. Philippe, M. Mani, P.F. Lenne, and T. Lecuit. 2018. Distinct contributions of tensile and shear stress on E-cadherin levels during morphogenesis. Nat Commun. 9:5021.

Kampmann, M., C.E. Atkinson, A.L. Mattheyses, and S.M. Simon. 2011. Mapping the orientation of nuclear pore proteins in living cells with polarized fluorescence microscopy. Nat Struct Mol Biol. 18:643–649.

Kenny, M., and I. Schoen. 2021. Violin SuperPlots: visualizing replicate heterogeneity in large data sets. Mol Biol Cell. 32:1333–1334.

Kim, S.A., C.Y. Tai, L.P. Mok, E.A. Mosser, and E.M. Schuman. 2011. Calcium-dependent dynamics of cadherin interactions at cell-cell junctions. Proc Natl Acad Sci U S A. 108:9857–9862.

Kostetskii, I., J. Li, Y. Xiong, R. Zhou, V.A. Ferrari, V.V. Patel, J.D. Molkentin, and G.L. Radice. 2005. Induced deletion of the N-cadherin gene in the heart leads to dissolution of the intercalated disc structure. Circulation research. 96:346–354.

le Duc, Q., Q. Shi, I. Blonk, A. Sonnenberg, N. Wang, D. Leckband, and J. de Rooij. 2010. Vinculin potentiates E-cadherin mechanosensing and is recruited to actin-anchored sites within adherens junctions in a myosin II-dependent manner. The Journal of Cell Biology. 189:1107–1115.

Leerberg, J.M., G.A. Gomez, S. Verma, E.J. Moussa, S.K. Wu, R. Priya, B.D. Hoffman, C. Grashoff, M.A. Schwartz, and A.S. Yap. 2014. Tension-sensitive actin assembly supports contractility at the epithelial zonula adherens. Curr Biol. 24:1689–1699.

Lewis, J.E., J.K. Wahl, 3rd, K.M. Sass, P.J. Jensen, K.R. Johnson, and M.J. Wheelock. 1997. Cross-talk between adherens junctions and desmosomes depends on plakoglobin. J Cell Biol. 136:919–934.

Li, Y., C.D. Merkel, X. Zeng, J.A. Heier, P.S. Cantrell, M. Sun, D.B. Stolz, S.C. Watkins, N.A. Yates, and A.V. Kwiatkowski. 2019. The N-cadherin interactome in primary cardiomyocytes as defined using quantitative proximity proteomics. J Cell Sci. 132.

Lord, S.J., K.B. Velle, R.D. Mullins, and L.K. Fritz-Laylin. 2020. SuperPlots: Communicating reproducibility and variability in cell biology. J Cell Biol. 219.

Mege, R.M., and N. Ishiyama. 2017. Integration of Cadherin Adhesion and Cytoskeleton at Adherens Junctions. Cold Spring Harbor perspectives in biology. 9.

Mehta, S.B., M. McQuilken, P.J. La Riviere, P. Occhipinti, A. Verma, R. Oldenbourg, A.S. Gladfelter, and T. Tani. 2016. Dissection of molecular assembly dynamics by tracking orientation and position of single molecules in live cells. Proc Natl Acad Sci U S A. 113:E6352–E6361.

Merkel, C.D., Y. Li, Q. Raza, D.B. Stolz, and A.V. Kwiatkowski. 2019. Vinculin anchors contractile actin to the cardiomyocyte adherens junction. Mol Biol Cell. 30:2639–2650.

Pang, S.M., S. Le, A.V. Kwiatkowski, and J. Yan. 2019. Mechanical stability of alphaT-catenin and its activation by force for vinculin binding. Mol Biol Cell. 30:1930–1937.

Radice, G.L., H. Rayburn, H. Matsunami, K.A. Knudsen, M. Takeichi, and R.O. Hynes. 1997. Developmental defects in mouse embryos lacking N-cadherin. Developmental biology. 181:64–78.

Seddiki, R., G. Narayana, P.O. Strale, H.E. Balcioglu, G. Peyret, M. Yao, A.P. Le, C. Teck Lim, J. Yan, B. Ladoux, and R.M. Mege. 2018. Force-dependent binding of vinculin to alpha-catenin regulates cell-cell contact stability and collective cell behavior. Mol Biol Cell. 29:380–388.

Shapiro, L., A.M. Fannon, P.D. Kwong, A. Thompson, M.S. Lehmann, G. Grubel, J.F. Legrand, J. Als-Nielsen, D.R. Colman, and W.A. Hendrickson. 1995. Structural basis of cell-cell adhesion by cadherins. Nature. 374:327–337.

Strale, P.O., L. Duchesne, G. Peyret, L. Montel, T. Nguyen, E. Png, R. Tampe, S. Troyanovsky, S. Henon, B. Ladoux, and R.M. Mege. 2015. The formation of ordered nanoclusters controls cadherin anchoring to actin and cell-cell contact fluidity. J Cell Biol. 210:1033.

Troyanovsky, R.B., A.P. Sergeeva, I. Indra, C.S. Chen, R. Kato, L. Shapiro, B. Honig, and S.M. Troyanovsky. 2021. Sorting of cadherin-catenin-associated proteins into individual clusters. Proc Natl Acad Sci U S A. 118.

Troyanovsky, S.M. 2023. Adherens junction: the ensemble of specialized cadherin clusters. Trends Cell Biol. 33:374–387.

Truong Quang, B.A., M. Mani, O. Markova, T. Lecuit, and P.F. Lenne. 2013. Principles of E-cadherin supramolecular organization in vivo. Curr Biol. 23:2197–2207.

Vermij, S.H., H. Abriel, and T.A. van Veen. 2017. Refining the molecular organization of the cardiac intercalated disc. Cardiovascular research. 113:259–275.

Vite, A., and G.L. Radice. 2014. N-cadherin/catenin complex as a master regulator of intercalated disc function. Cell communication & adhesion. 21:169–179.

Wu, Y., X. Jin, O. Harrison, L. Shapiro, B.H. Honig, and A. Ben-Shaul. 2010. Cooperativity between trans and cis interactions in cadherin-mediated junction formation. Proc Natl Acad Sci U S A. 107:17592–17597.

Wu, Y., J. Vendome, L. Shapiro, A. Ben-Shaul, and B. Honig. 2011. Transforming binding affinities from three dimensions to two with application to cadherin clustering. Nature. 475:510–513.

Yao, M., W. Qiu, R. Liu, A.K. Efremov, P. Cong, R. Seddiki, M. Payre, C.T. Lim, B. Ladoux, R.M. Mege, and J. Yan. 2014. Force-dependent conformational switch of alpha-catenin controls vinculin binding. Nat Commun. 5:4525.

Yonemura, S., Y. Wada, T. Watanabe, A. Nagafuchi, and M. Shibata. 2010. alpha-Catenin as a tension transducer that induces adherens junction development. Nat Cell Biol. 12:533–542.

